# Rme-6 integrates EGFR trafficking and signalling to regulate ERK1/2 signalosome dynamics

**DOI:** 10.1101/2023.05.05.539436

**Authors:** Fahad Alshahrani, Zhou Zhu, Filipe Ferreira, Alasdair McComb, Hannes Maib, Sara Pruzina, Darren Robinson, David Murray, Elizabeth Smythe

## Abstract

Epidermal growth factor receptor (EGFR) signalling results in a variety of cell behaviours, including cell proliferation, migration and apoptosis, which depend on cell context. Here we have explored how the Rab5GEF, Rme-6, regulates EGFR signalling by modulating endocytic flux. We demonstrate that Rme-6, which acts early in the endocytic pathway, regulates EGFR trafficking through an endocytic compartment that is competent for ERK1/2 signalling. While overexpression of Rme-6 results in enhanced ERK1/2 nuclear localisation and c-Fos activation, loss of Rme-6 results in aberrant ERK1/2 signalling with increased cytoplasmic ERK1/2 phosphorylation (Thr202/Tyr204) but decreased ERK1/2 nuclear translocation and c-Fos activation, the latter leading to decreased cell proliferation. Phosphorylation of ERK1/2 by protein kinase 2 (CK2) is required for its nuclear translocation and our data support a model whereby Rme-6 provides a scaffold for a population of CK2 which is required for efficient nuclear translocation of ERK1/2. Rme-6 is itself a substrate for CK2 on Thr642 and Ser996 and phosphorylation on these sites can activate its Rab5GEF activity and endocytic trafficking of EGFR. Together our results indicate that Rme-6 co-ordinates EGFR trafficking and signalling to regulate the assembly and disassembly of an ERK1/2 signalosome.

**Summary statement:** Here we demonstrate how Rme-6, a Rab5GEF, co-ordinates trafficking and signalling of EGFR on the early endocytic pathway to ensure appropriate regulation of downstream ERK1/2 signalling.

## Introduction

Ligand activation of receptor tyrosine kinases, such as epidermal growth factor receptor (EGFR), gives rise to a variety of downstream cell behaviours including cell proliferation, migration apoptosis and differentiation. These downstream signalling cascades include Ras activated ERK1/2 and PI3Kinase activation of Akt (Lemmon and Schlessinger, 2010; Schlessinger, 2014). How cells respond to EGFR activation depends on their location and thus cell-context dependent regulation of signalling needs to be tightly regulated. A key question is how cells in a particular environment interpret specific cues. One mechanism that allows such a diversity of signalling outcomes is signal regulation by endocytosis (Moore et al., 2018; Sigismund et al., 2021; Sorkin and von Zastrow, 2009). Both *in vitro* and *in vivo* studies have demonstrated that following activation at the cell surface, receptors such as EGFR continue to signal as they traverse the endocytic pathway and, importantly, signalling output is both qualitatively as well as quantitatively different depending on the location of the activated receptor in the endocytic pathway (Caldieri et al., 2018; Eden et al., 2010; Sapmaz and Erson-Bensan, 2023; Sigismund et al., 2008; Vieira et al., 1996). Delivery of receptors to lysosomes can attenuate signalling through receptor degradation while membrane-associated signalosomes can qualitatively regulate signalling. Signalosomes are specialised membrane domains harbouring specific scaffolds, adaptors, kinases and phosphatases. Such segregation is thought to increase both the strength and specificity of signalling (Kholodenko, 2009). Precedents for endocytic signalosomes are the p14 scaffold for MAPK signalling on late endosomes (Teis et al., 2002), endocytosis-dependent Wnt signalosomes (Colozza and Koo, 2021), clathrin coated lattices acting as signalling hubs (Cabral-Dias et al., 2022), while for EGFR it is well-established that signalling occurs from the plasma membrane as well as endosomes (Tomas et al., 2014; Villasenor et al., 2015; Wang et al., 2002). It follows that the rate at which the receptor moves along the endocytic pathway through different signalosomes, its flux, will be key to determining signalling outputs. The activity of individual components of the endocytic machinery is essential to regulate endocytic flux of receptors. Cell fate, for example, can be altered by modulating levels of the endosomal fusion machinery, to prolong the residence of receptors in specific endosomal compartments (Villasenor et al., 2015).

The Rab family of small GTPases are key regulators of membrane traffic which cycle between an inactive GDP bound conformation in the cytoplasm and an active GTP form which is membrane-associated. Conversion from the GDP to the GTP form is mediated by guanine nucleotide exchange factors (GEFs). Individual Rabs are found localised to specific intracellular membranes (Homma et al., 2021; Lamber et al., 2019; Muller and Goody, 2018). Rab5 is localised to the early endocytic pathway where it regulates multiple aspects of endosomal function, including clathrin coated vesicle uncoating, endosomal fusion, motility and signalling (McLauchlan et al., 1998; Mottola et al., 2010; Shin et al., 2005; Simonsen et al., 1998; Stenmark, 2009; Zeigerer et al., 2012). Rab5 interacts with greater than 20 different cytosolic proteins (effectors) and it recruits subsets of these proteins to perform its diverse functions (Christoforidis et al., 1999; Miaczynska et al., 2004 ; Nielsen et al., 2000; Schnatwinkel et al., 2004). In vivo the Rab5 effector APPL1 is required for a subset of Akt signalling (Schenck et al., 2008).

The recruitment of a given complement of Rab effectors for a particular function is driven by GEFs which activate Rabs on specific membranes (Blumer et al., 2013; Semerdjieva et al., 2008) Rme-6, also known as GAPex5 and GAPVD1, is a Rab5GEF which was originally identified in the function of clathrin coated vesicles at the plasma membrane in *C.elegans* (Sato et al., 2005). Our subsequent studies showed that it regulates the uncoating of the AP2 adaptor complex from clathrin coated vesicles in mammalian cells (Semerdjieva et al., 2008). In addition to its Rab5GEF (Vps9) domain, it has a RasGAP domain which has been shown to promote GTP hydrolysis on Ras in vitro, thus inactivating it (Su et al., 2007).

Here we have investigated the role of Rme-6 in establishing a Rab domain specialised for EGFR signalling, a signalosome, using HeLa cells and MCF10A cells which are a non-tumorigenic immortalised epithelial cell line (Vale et al., 2021). We find that Rme-6 integrates EGFR trafficking and signalling in these cell lines, acting in the regulated assembly and disassembly of an EGFR signalosome specific for ERK1/2 signalling and thus ensuring appropriate downstream cell behaviour.

## Results

### Loss of Rme-6 affects EGFR signalling in HeLa cells

As a first step to understand whether Rme-6 might integrate endocytic trafficking and signalling, we generated HeLa cell lines lacking Rme-6, Rme-6^-ve^, using CRISPR technology (Figure 1a). Using Alexa^555^EGF as a proxy for EGFR internalisation we found that initial rates of uptake were identical in HeLa and Rme-6^-ve^ cells (Figure 1b). We then measured rates of EGFR degradation of EGFR in HeLa and Rme6^-ve^ cell lines. We found that the rate of EGFR degradation in response to incubation with EGF was delayed in Rme6^-ve^ cells (Figure 1c and d) suggesting that, in the absence of Rme-6, EGFR is not efficiently delivered to lysosomes. In support of delayed receptor degradation, we found that overall levels of EGFR, measured by Western blotting, are increased in Rme-6^-ve^ cells compared with HeLa cells (Figure 1e). Cell surface levels of EGFR, measured by flow cytometry, were decreased in Rme-6^-ve^ cells, suggesting that in the absence of Rme-6 EGFR accumulates intracellularly (Figure 1f). We observed a similar increase in overall expression of EGFR and decreased surface expression in MCF10A cells where Rme-6 levels had been reduced by siRNA (Supplementary Figure 1a-c).

**Figure 1:**
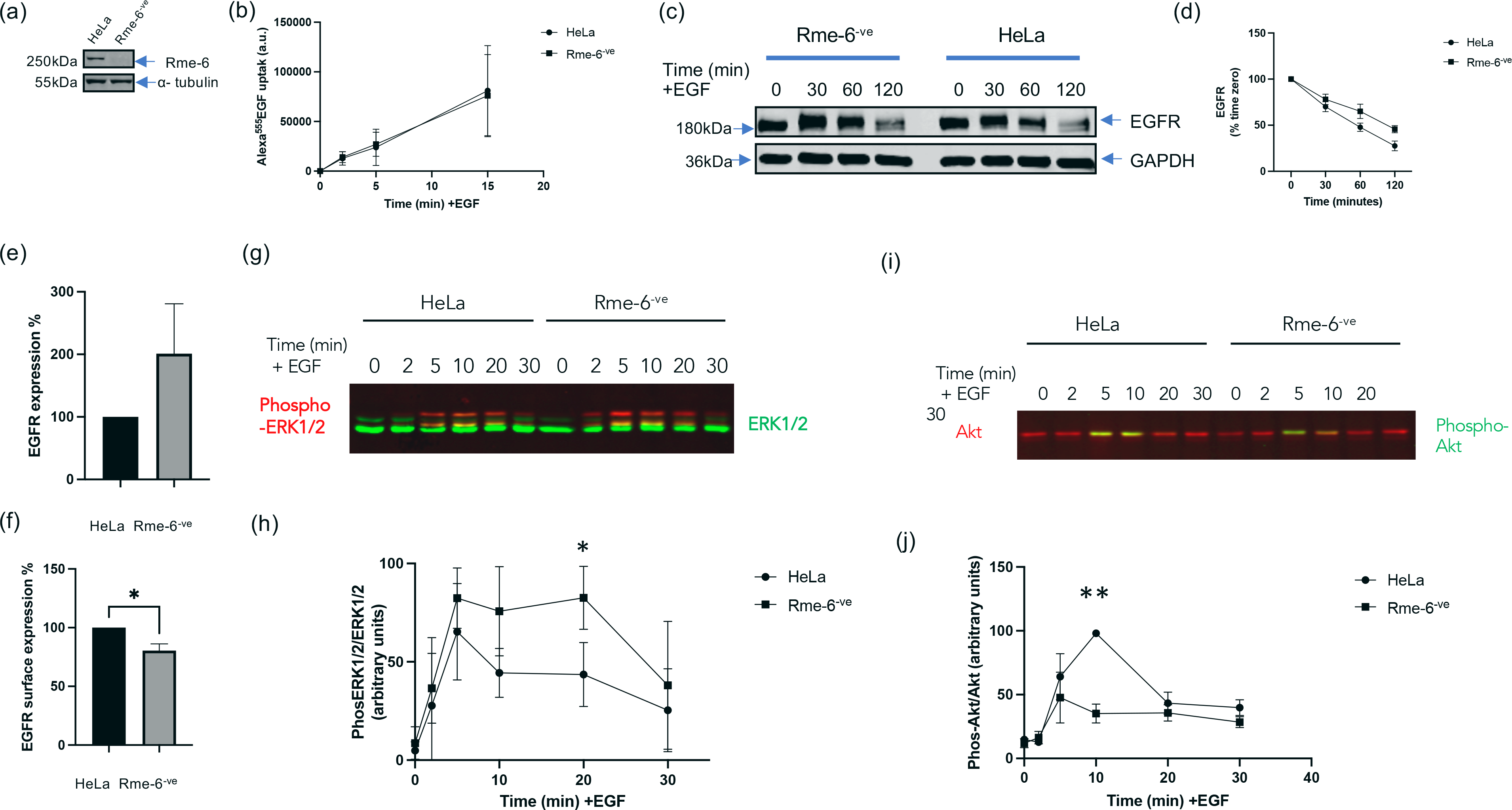
Rme-6 qualitatively affects EGFR signalling in HeLa cells. (a) Western blots of lysates from HeLa cells and HeLa cells where Rme-6 has been knocked out by CRISPR (Rme-6^-ve^). Blots were probed with antibodies against Rme-6 and α-Tubulin as a loading control. (b) HeLa and Rme-6^-ve^ cells were incubated with Alexa^555^EGF for the indicated times. Following acid-strip, Alexa^555^EGF fluorescence was measured and quantified using ImageJ. Data are the mean +/-SEM of three independent experiments. (c) Representative Western blots of lysates from HeLa and Rme-6^-ve^ cells following treatment with EGF (5ng/ml) for the indicated times. Blots were probed with antibodies against EGFR and GAPDH as a loading control. (d) Quantitation of EGFR degradation from 3 independent experiments. Data are expressed as mean +/- SEM. (e) Expression of EGFR in HeLa and Rme-6^-ve^ cells determined by Western blotting. Data are the mean +/-SEM. (f) Surface expression of EGFR in HeLa and Rme-6^-ve^ cells was measured by Facs. Data are the mean +/-SEM of three independent experiments. *:p<0.05 (g) Representative Western blot showing total ERK1/2 (green) and phosphorylated ERK1/2 (red) in HeLa and Rme-6^-ve^ cells following EGF stimulation (5ng/ml) for the indicated times. (h) Quantitation of the ratio of phospho-ERK1/2 to total ERK1/2 from three independent experiments. Data are expressed as mean +/- SEM. *<0.05 (i) Representative Western blot showing total Akt (red) and phosphorylated Akt (green) in HeLa and Rme-6^-ve^ cells following EGF stimulation (5ng/ml) for the indicated times. (j) Quantitation of the ratio of phosphorylated Akt to total Akt from three independent experiments. Data are expressed as mean +/- SEM. **: p<0.01

In HeLa cells EGF-dependent EGFR activation results in ERK1/2 phosphorylation on Thr202/Tyr204, which can be detected by Western blotting using phospho-specific antibodies (Roskoski, 2012). Treatment with EGF (5ng/ml) increases levels of Thr202/Tyr204 phosphorylation which peak at approximately 5 minutes and then slowly decline (Figure 1g and h). In the absence of Rme-6, ERK1/2 signalling is both enhanced with increased overall levels of Thr202/Tyr204 phosphorylation which are sustained until 20 minutes before declining (Figure 1g and h). EGFR activation also results in increased Akt phosphorylation (Ser473) with a peak at 5 minutes which then declines over the next 25 minutes (Figure 1i and j). In contrast to ERK1/2 signalling, loss of Rme-6 results in a reduction in the extent of Akt signalling at all time points measured (Figure 1i and j). In MCF10A cells, siRNA-mediated knockdown of Rme-6 increased expression levels of both ERK1/2 and Akt resulting in overall increased levels of both phospho-ERK1/2 and phospho-Akt (Supplementary Figure 1d-f). Consistent with delayed EGFR degradation leading to prolonged signalling, we also found a trend that Ras activation following EGF stimulation was prolonged in both HeLa and MCF10A cells (Supplementary Figure 2 a-d).

### Loss of Rme-6 affects EGFR trafficking

We next wanted to understand whether changes in EGFR trafficking in the absence of Rme-6 were leading to the observed changes in EGFR signalling, as well as causing its intracellular accumulation and reduced cell surface expression. We therefore measured the extent of Alexa^555^EGF (5ng/ml) co-localisation with APPL1 and EEA1 positive compartments, following incubation for 5 and 15 minutes. In control HeLa cells, Alexa^555^EGF colocalises with APPL1 early (∼5 min) after internalisation as has been previously observed (Miaczynska et al., 2004). By contrast in Rme-6^-ve^ cells, the degree of overlap between Alexa^555^EGF and APPL1 was significantly higher than control cells after both 5 and 15 minutes (Figure 2a and b), indicating that loss of Rme-6 prolongs residence of EGFR in APPL1- positive compartments. Conversely, delivery of Alexa^555^EGF to EEA1 positive endosomes, which are thought to be reached at later times post internalisation (Miaczynska et al., 2004), was significantly delayed in Rme-6^-ve^ cells compared to control HeLa cells. Together this suggests that Rme-6 is required for exit from APPL1 positive endosomes.

**Figure 2:**
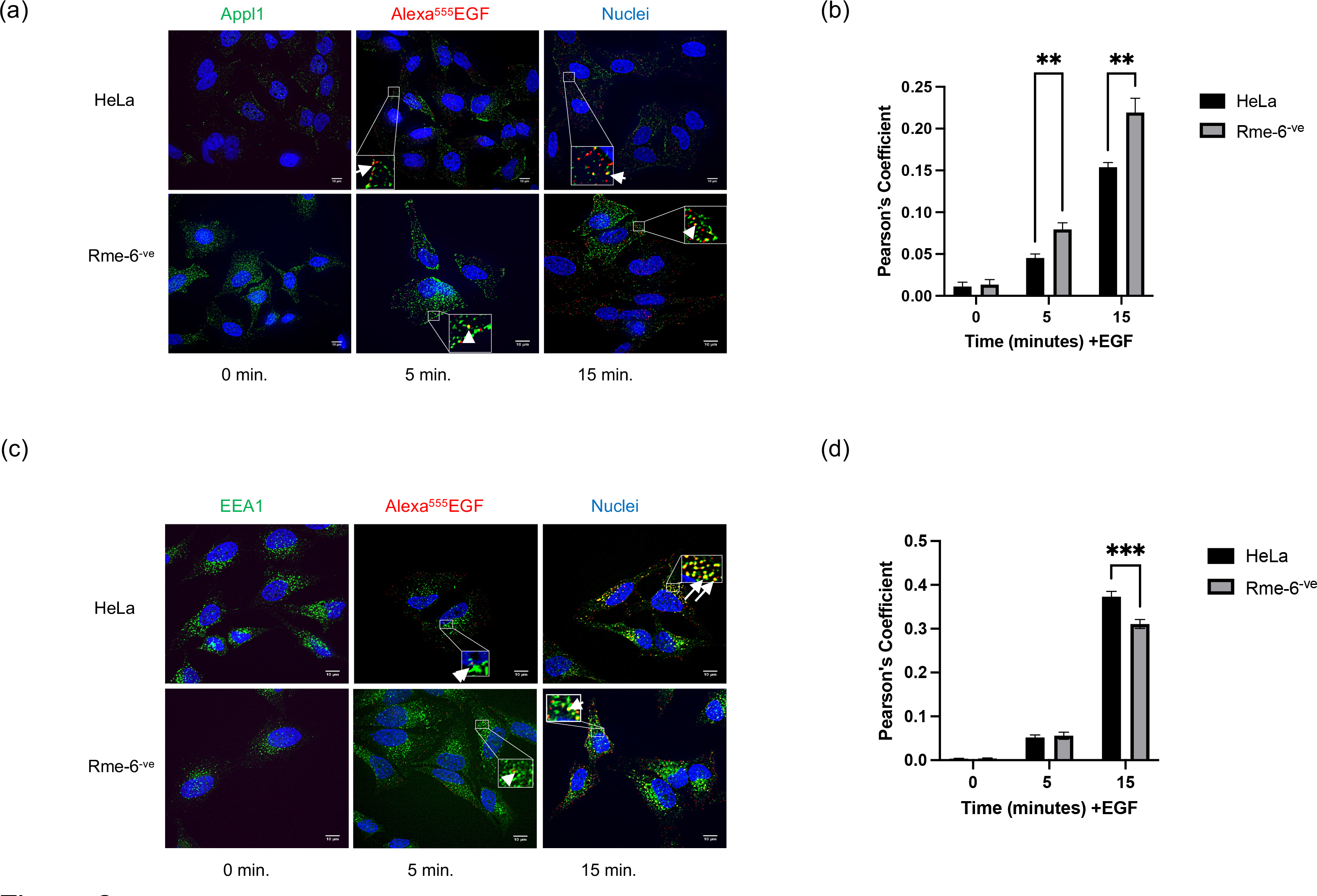
Loss of Rme-6 affects EGFR trafficking in HeLa cells. (a) Following serum starvation, HeLa and Rme-6^-ve^ cells were incubated with Alexa^555^ EGF (5ng/ml) for the indicated times. Cells were fixed and stained for APPL1 (green) and nuclei stained with Dapi (blue). The white boxes are magnified images with examples of overlap between Alexa^555^EGF (red) and APPL1(green) indicated with white arrows. Bar: 10µM (b) The extent of overlap between APPL1 and Alexa^555^EGF was determined by Pearson’s Coefficient using Image J. Data are the mean +/- SEM of 3 independent experiments where at least 30 cells were analysed per condition. **: p<0.01; (c) Following serum starvation, HeLa and Rme-6^-ve^ cells were incubated with Alexa^555^EGF (5ng/ml) for the indicated times. Cells were fixed and stained for EEA1 (green) and the nuclei stained in blue. The white boxes are magnified images with examples of overlap between Alexa^555^EGF (red) and EEA1 (green) indicated with white arrows. Bar: 10µM (d) The extent of overlap between EEA1 and Alexa^555^EGF was determined by Pearson’s Coefficient using Image J. Data are the mean +/- SEM of 3 independent experiments where at least 30 cells were analysed per condition. ***: P<.0001.

### CK2 phosphorylation regulates Rme-6

We next wanted to explore how Rme-6 might regulate the transit of EGFR through endocytic compartments. Rme-6 has many different phosphorylation sites (Ducommun et al., 2019; Guillen et al., 2020; Ibrahim et al., 2021) and our previous studies had implicated phosphorylation of Rme-6 in the regulation of its activity (Semerdjieva et al., 2008). We have shown that Rme-6 acts early (2- 5min) in the endocytic pathway following uncoating of clathrin coated vesicles (Semerdjieva et al., 2008). Our observations that the absence of Rme-6 changes the co-localisation of Alexa^555^EGF with both APPL1 and EEA1 (Figure 2 a-d) also supports a role for Rme-6 soon after internalisation. We were therefore particularly interested in the role that clathrin coated vesicle kinases might play in Rme-6 function and found that a population of CK2, a major clathrin coated vesicle kinase (Korolchuk and Banting, 2002), is associated with Rme-6 when it is immunoprecipitated from mammalian cells (Supplementary Figure 3 a and b). Analysis of the sequence of Rme-6 identified several phosphorylation sites (Net phos 2.0) and, in particular, we explored the roles of Ser996 and Thr642 as likely CK2 consensus phosphorylation sites. Using baculoviral expression we purified Rme6^wt^, a non-phosphorylatable version, Rme-6^T642A/S996A^, and a phospho-mimetic version, Rme-6^T642D/S996D^, and confirmed that while Rme-6 ^wt^ was efficiently phosphorylated by CK2 in vitro, there was significantly reduced CK2 phosphorylation of both the non-phosphorylatable Rme-6^T642A/S996A^ and the phospho-mimetic form, Rme-6^T642D/S996D^ (Figure 3a and b). This supports these two sites as key in vitro CK2 substrates in Rme-6. The low level of CK2 phosphorylation observed in Rme-6^T642A/S996A^ and Rme-6^T642D/S996D^ is likely due to phosphorylation of other CK2 sites in Rme-6.

**Figure 3:**
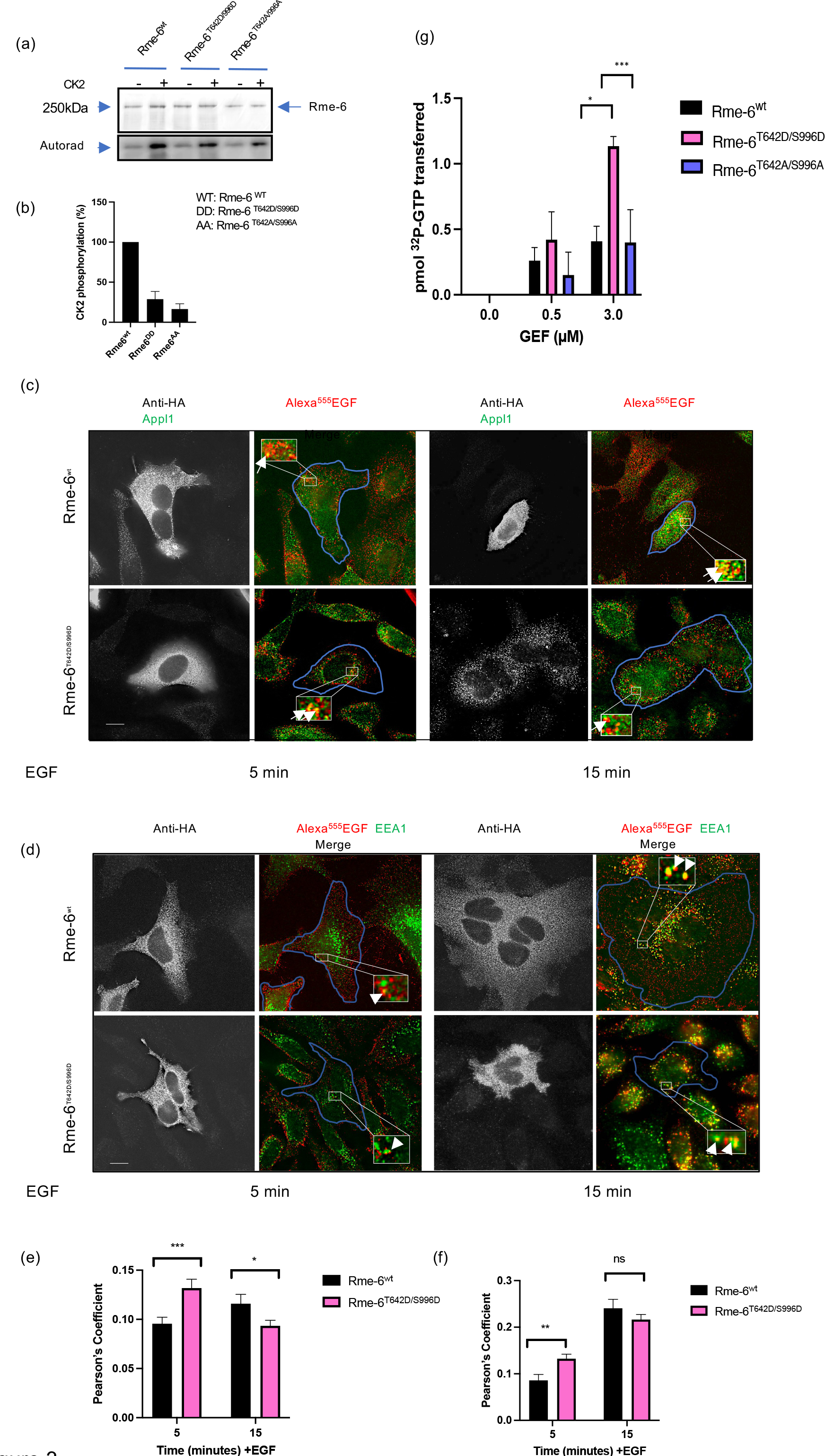
CK2 regulates Rme-6 activity. (a) *In vitro* kinase assay using γ^32P^-ATP with purified Rme-6^wt^, Rme-6^T642A/S996A^ and Rme-6^T642D/S996D^ (0.1µM) in the presence or absence of CK2 as indicated. Top panel shows SDS gel of purified proteins with the corresponding autoradiograph below. (b) Quantitation of phosphorylation of Rme-6^wt^, Rme-6^T642A/S996A^ and Rme-6^T642D/S996D^ by CK2 expressed as a percentage where phosphorylation of Rme-6^wt^ has been set at 100%. Data represent mean +/- SEM of 3 independent experiments. (c) Fluorescence images of HeLa cells transfected with HA-tagged Rme-6^wt^ and Rme-6^T642D/S996D^ (grey images in left panels, transfected cells highlighted in blue in right hand panel) which were incubated with Alexa^555^EGF (5ng/ml) for the indicated times and then fixed and stained with anti-APPL1 antibodies (green). Enlarged inserts show overlap between APPL1 and Alexa^555^EGF indicated by arrows. Bar: 10µM (d) Fluorescence images of HeLa cells transfected with HA-tagged Rme-6^wt^ and Rme-6^T642D/S996D^ (grey images in left panels, transfected cells highlighted in blue in right hand panel) which were incubated with Alexa^555^EGF (5ng/ml) for the indicated times and then fixed and stained with anti-EEA1 antibodies (green). Enlarged inserts show co-localisation between EEA1 and Alexa^555^EGF indicated by white arrows. Bar: 10µM (e) Quantitation of the overlap of Alexa^555^EGF with APPL1. Results are expressed as the mean +/- SEM from three independent experiments where between 15 and 25 cells cells were measured in each experiment. *: p<0.05; ***: P<0.01. (f) Quantitation of the overlap of Alexa^555^EGF with EEA1. Results are expressed as the mean +/- SEM of data from three independent experiments where between 14 and 19 cells were measured in each experiment. **: p<0.05, ns: not significant. (g) Purified Rme-6^wt^, Rme-6^T642A/S996A^ and Rme-6^T642D/S996D^ were incubated with GST-Rab5GDP and α^32P^-GTP and the amount of GTP exchange measured after 30 min at 30°C. Values are expressed as mean +/- SEM of 3 independent experiments each performed in duplicate. *: p<0.05; ***: p<0.001

Following these *in vitro* experiments, we asked whether phosphorylation on these two sites affected endocytic flux of EGFR through the early endocytic pathway. We thus transfected HeLa cells with HA-tagged Rme6^wt^ and Rme-6^T642D/S996D^ and, following incubation with Alexa^555^EGF (5ng/ml) for 5 and 15 minutes, we measured the co-localisation of EGF with APPL1- and EEA1-positive compartments.

We found that compared to cells overexpressing Rme-6^wt^, there is significantly more co-localisation of EGF with APPL1 at 5 minutes in the presence of overexpressed Rme-6^T642D/S996D^. By contrast at 15 mins there is significantly less overlap between Alexa^555^EGF following overexpression of Rme- 6^T642D/S996D^ compared to overexpression of Rme-6^wt^ (Figure 3c and e). Overexpression of Rme-6^T642D/S996D^ resulted in significantly higher overlap between Alexa^555^EGF and EEA1 at 5 minutes compared to overexpression of Rme-6^wt^ (Figure 3 d and f). Together these results suggest that Rme-6^wt^ may have a dominant negative effect over the endogenous Rme-6 protein resulting in prolonged residence of activated EGFR in an APPL1 compartment. The phospho-mimetic Rme-6^T642D/S996D^ appears to enhance delivery through an APPL1 compartment to an EEA1 positive endosome.

Combined with the results of the *in vitro* kinase assay, these data suggested that phosphorylation by CK2 on Ser996 and Thr642 is necessary for the Rme-6 dependent movement of EGFR through APPL1- and EEA1-positive compartments. Since this would likely require the Rab5GEF activity of Rme-6, we carried out guanine nucleotide exchange assays on Rab5 using purified Rme6^wt^, Rme-6^T642A/S996A^ and Rme-6^T642D/S996D^. Consistent with our previous observation that isolated Rme-6 is auto-inhibited (Semerdjieva et al., 2008), we found that both Rme6^wt^ and Rme-6^T642A/S996A^ showed no detectable activity in *in vitro* rab5GEF assays whereas Rme-6^T642D/S996D^ was able to catalyse GTP exchange in the absence of any Rme-6 binding partners (Figure 3g). Together these results support a model where CK2 phosphorylation alters the conformation of Rme-6 resulting in Rab5GEF activation and progression of EGFR along the endocytic pathway to APPL1- and EEA1-positive compartments.

### Rme-6 is essential for efficient nuclear translocation of ERK1/2

In order to regulate downstream cell behaviours, ERK1/2 needs to enter the nucleus where it activates early genes including c-Fos (Murphy and Blenis, 2006). Previous studies have shown that CK2-mediated ERK1/2 phosphorylation on Ser 244 and 246 facilitates its binding to importin7 which in turn allows its nuclear translocation (Schevzov et al., 2015). Since we had shown that there is a population of CK2 associated with Rme-6 which appears to regulate its Rab5GEF activity, we were curious as to whether CK2 associated with Rme-6 might also be important for ERK1/2 nuclear translocation. To address this question, we measured nuclear accumulation of phospho-ERK1/2 by immunofluorescence. We first confirmed that treatment with the CK2 inhibitor, TBB, inhibited nuclear translocation of phospho-ERK1/2 in HeLa cells (Plotnikov et al., 2019) (Supplementary Figure 3c). We then measured nuclear accumulation of phospho-ERK1/2 in MCF10A cells, treated either with control siRNA or siRNA targeting Rme-6, and incubated with EGF (5ng/ml). Surprisingly, we found that MCF10A cells where Rme-6 had been knocked down by siRNA, showed reduced ERK1/2 nuclear fluorescence compared to cells treated with control siRNA (Figure 4 a and b). Once in the nucleus, ERK1/2 phosphorylates many substrates, one of which is the transcription factor c-Fos, which is essential for subsequent proliferation (Murphy and Blenis, 2006). Consistent with the reduced nuclear ERK1/2 observed when Rme-6 levels are lowered, we also observed lower levels of phospho-c-Fos in MCF10A cells when Rme-6 levels were reduced by siRNA (Figure 4c and d).

**Figure 4:**
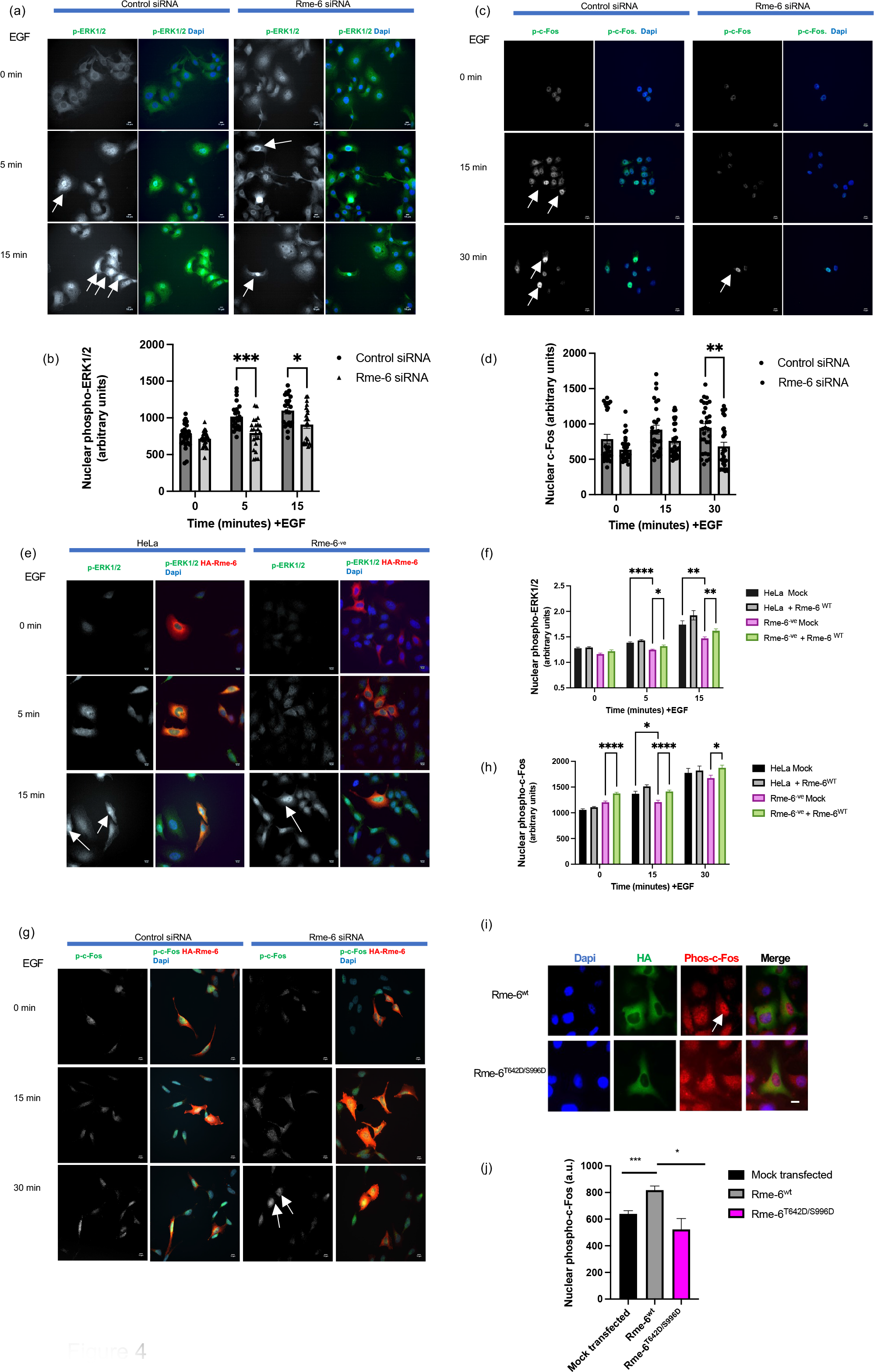
Rme-6 ensures efficient ERK1/2 nuclear translocation and c-Fos phosphorylation. (a) MCF10A cells treated with control siRNA or siRNA targeting Rme-6 were incubated with 5ng/ml EGF for the indicated times and stained for phospho-ERK1/2 (green). Nuclei were stained with Dapi (blue). White arrows indicate increased phospho-ERK1/2 in the nucleus. Scale bar: 10µM (b) Quantitation of nuclear localisation of phospho-ERK1/2 in three independent experiments. Data are expressed as mean +/-SEM. *: p<0.05; ***: p<0.001 (c) Images of MCF10A cells treated with control siRNA or siRNA targeting Rme-6 which were incubated with 5ng/ml EGF for the indicated times and stained for phospho-c-Fos (green). Nuclei were stained with Dapi (blue). White arrows indicate increased phospho-c-Fos in the nucleus. Scale bar: 10µM (d) Quantitation of the levels of nuclear phospho-c-Fos in three independent experiments. Data are expressed as mean +/-SEM. **: p<0.01 (e) HeLa and Rme-6^-ve^ cells were were transfected with Rme-6^wt^ (red) and incubated with 5ng/ml EGF for the indicated times. Cells were fixed and stained for phospho-ERK1/2 (green). Nuclei were stained with Dapi (blue). White arrows indicate a cell with increased phospho-ERK1/2 in the nucleus. Scale bar: 10µM (f) Quantitation of nuclear localisation of phospho-ERK1/2 in two independent experiments. Data are expressed as mean +/-SD. *: p<0.05; **: p<0.01; ****: P<0.0001 (g) HeLa and Rme-6^-ve^ cells were transfected with Rme-6^wt^ (red) and incubated with 5ng/ml EGF for the indicated times. Cells were fixed and stained for phospho-c-Fos (green). Nuclei were stained with Dapi (blue). White arrows indicate a cell with increased phospho-c-Fos in the nucleus. Scale bar: 10µM (h) Quantitation of nuclear localisation of phospho-c-Fos in two independent experiments. Data are expressed as mean +/-SD. *: p<0.05; ****: p<0.001 (i) HeLa cells were transfected with HA-tagged Rme-6^wt^ or Rme-6^T642D/S996D^ and incubated with 5ng/ml EGF for 30 min before being fixed and stained with antibodies to HA (green) and phospho-c-Fos (red). Nuclei were stained with Dapi (blue). White arrow indicates increased levels of nuclear phospho-c-Fos in a cell overexpressing Rme-6^wt^. Scale bar: 10µM (j) Quantification of nuclear phospho-c-Fos in mock transfected HeLa cells and HeLa cells transfected with Rme-6^wt^ or Rme-6^T642D/S996D^. Data are expressed as the mean +/- SEM of at least 5 independent experiments where 15-25 cells were analysed per experiment. Significance was determined with a two-tailed Students T-test. *: p<0.05; ***: p<0.001.

To confirm that the reduced levels of nuclear ERK1/2 and phospho-c-Fos resulted specifically from a loss of Rme-6, we carried out rescue experiments in HeLa and Rme-6^-ve^ cell lines. Cells were transfected with HA-tagged Rme-6^wt^ and the levels of nuclear ERK1/2 measured following incubation with EGF. Similar to our results in MCF10A cells, we observed that Rme-6^-ve^ cells showed less nuclear accumulation of ERK1/2 compared to control HeLa cells (Figure 4 e and f). Transfection of HA-Rme-6^wt^ resulted in increased nuclear ERK1/2 in control HeLa cells suggesting that increasing levels of Rme-6 could promote ERK1/2 translocation. In Rme-6^-ve^ cells transfection of HA-Rme-6^wt^ resulted in some rescue of nuclear ERK1/2 although not to the levels seen in control HeLa cells (Figure 4 e and f). We also measured the levels of phospho-c-Fos in HeLa and Rme-6^-ve^ cells. As we had observed in MCF10A cells, loss of Rme-6 resulted in reduced levels of phospho-c-Fos (Figure 4 g and h).

Expression of HA-Rme-6^wt^ was however sufficient to rescue phospho-c-Fos levels in Rme-6^-ve^ cells to those of HeLa cells, suggesting that the levels of ERK1/2 translocated into the nucleus following overexpression of HA-Rme-6^wt^ were sufficient for downstream signalling.

We had shown that Rme-6^T642D/S996D^ could rescue the trafficking defects observed following overexpression of Rme6^wt^ (Figure 3 c-f). If there is an Rme-6 mediated coupling between EGFR trafficking and signalling it would be predicted that CK2 phosphorylation of Rme-6 might also impact its signalling. To test this possibility, we measured the levels of phosphorylated c-Fos in the nucleus following overexpression of Rme-6^wt^ and Rme-6^T642D/S996D^. We found that overexpression of Rme-6^wt^ significantly increased levels of nuclear phospho-c-Fos compared to mock transfected controls. By contrast expression of Rme-6^T642D/S996D^ resulted in levels of nuclear c-Fos similar to those observed in mock transfected cells (Figure 3 i and j). Thus the phospho-mimetic Rme-6^T642D/S996D^ rescues the dominant negative effect of Rme-6 overexpression on signalling as well as trafficking.

### Loss of Rme-6 reduces cell proliferation and increases apoptosis in MCF10A cells

Our results suggest that Rme-6 integrates EGFR trafficking and signalling. Therefore we would expect to see effects on downstream cell behaviour when Rme-6 levels are modulated. We observed when we treated MCF10A cells with siRNA targeting Rme-6 that there was a dramatic decrease in cell number compared to cells treated with control siRNA (Figure 5a). We wanted to test whether this was due to cell proliferation and/or apoptosis. We first used BrdU incorporation and FACS analysis to determine whether loss of Rme-6 affects the length of S phase in MCF10A cells. We found that compared to cells treated with control siRNA, MCF10A cells treated with siRNA targeting Rme-6 showed a significantly reduced number of cells in S phase with a corresponding increase in the number of cells in G1 (Figure 5b), indicating that Rme-6 dependent changes in ERK1/2 signalling correlate with changes in cell proliferation. To test whether the reduction in cell number could also be due to apoptosis we measured Caspase3 expression in MCF10A cells following siRNA mediated knockdown of Rme-6. We found that there was a significant increase in Caspase3 levels in cells with reduced Rme-6, indicative of increased apoptosis (Cohen, 1997) (Figure 5c and d). This correlated with the increased activated Akt signalling observed in MCF10A cells following loss of Rme-6 (Supplementary Figure 3c-e).

**Figure 5:**
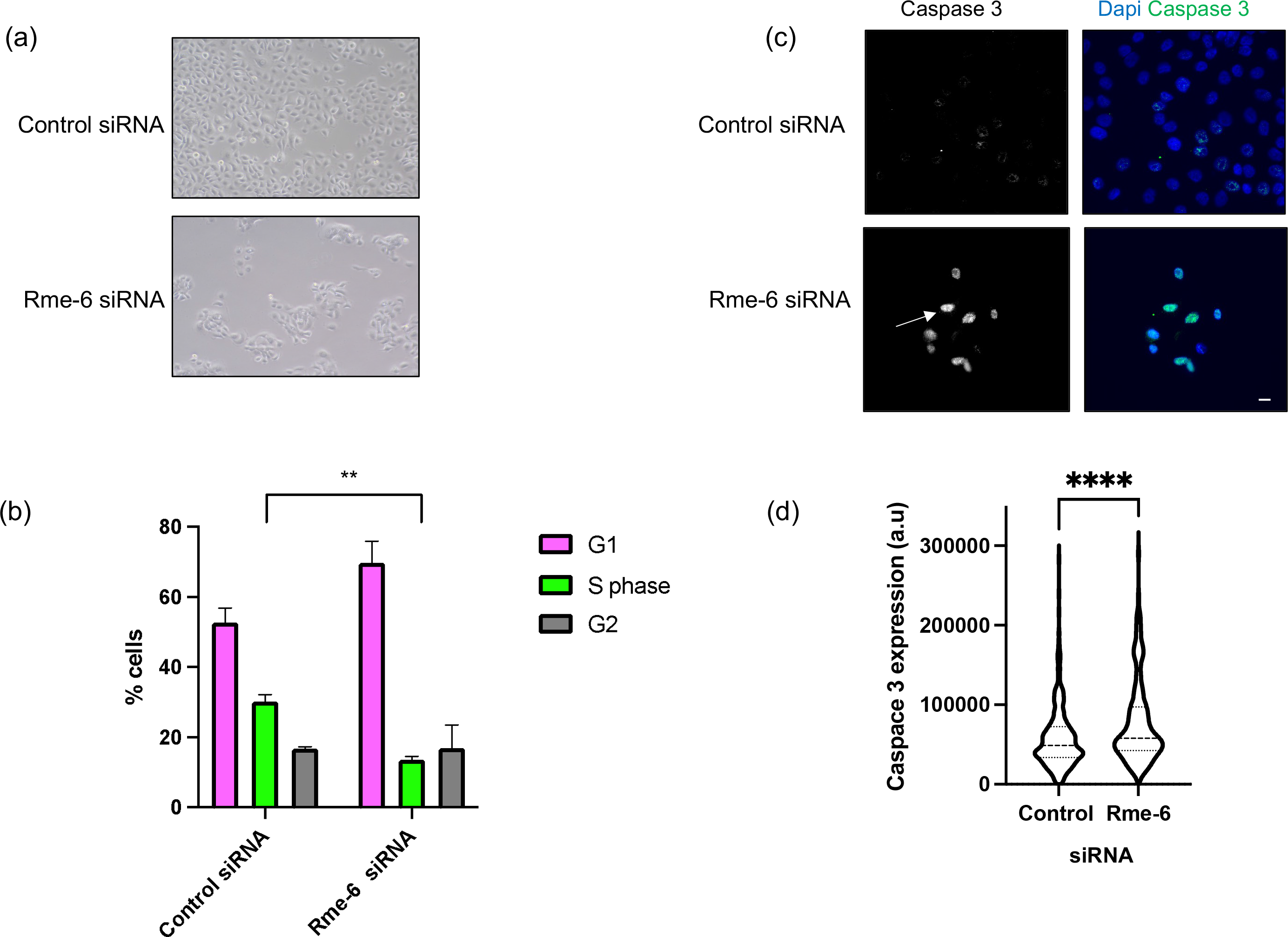
Loss of Rme-6 affects cell behaviour. (a) Equal numbers of MCF10A cells treated with control siRNA or siRNA targeting Rme-6 were seeded and analysed by brightfield after 5 days. (b) MCF10A cells treated with control siRNA or siRNA targeting Rme-6 were labelled with BrdU to estimate the length of S-phase. Data are from three independent experiments and are expressed as mean +/-SEM. **: p<0.01 (c) MCF10A cells treated with control siRNA or siRNA targeting Rme-6 were fixed and labelled with anti-Caspase3 antibodies (green), nuclei were labelled with DAPI in blue. The white arrow indicates an example of increased Caspase3 in the nucleus of a cell treated with siRNA against Rme-6. Bar: 10µM. (d) Quantification of Caspase3 labelling from 3 independent experiments where >25 cells were analysed in each experiment. Data are expressed as mean +/-SEM; Significance was determined using a Student T-Test, ****: p<.0001.

## Discussion

The aim of our study was to investigate whether the Rab5GEF, Rme-6, could integrate signalling and trafficking of activated EGFR. We show that Rme-6 co-ordinates EGFR trafficking and signalling to establish a signalosome which is required for effective ERK1/2 signalling outcomes, including cell proliferation and apoptosis. In the process we have gained considerable mechanistic insight as to how the Rme-6 driven ERK1/2 signalosome is assembled and disassembled. Such co-ordination between receptor trafficking and signalling leads to appropriate ERK1/2 signalling.

### Assembly and disassembly of an Rme-6 driven ERK1/2 signalosome

We explored the function of Rme-6 in HeLa cells where Rme-6 had been knocked out by CRISPR, as well as in MCF10A cells where Rme-6 levels had been reduced by siRNA. We found that in HeLa cells loss of Rme-6 altered the trafficking of EGFR through the early endosomal system, causing increased association with APPL1 endosomes and a delay in reaching EEA1 positive compartments, consistent with a longer residence time in a compartment reached early after internalisation. Overexpression of Rme-6^wt^ caused a similar delay in trafficking of EGFR to early endosome compartments indicating that overexpression causes dominant negative effects on Rme-6 function which could be relieved by CK2 phosphorylation (see below). The delay in the trafficking of EGFR in the absence of Rme-6 correlated with alterations in signalling. Loss of Rme-6 resulted in increased phosphorylation of ERK1/2^Thr202/Tyr204^ which we attribute to prolonged EGFR residence time in a compartment competent for EGFR dependent activation of ERK1/2 on Thr 202/Tyr204. Surprisingly, however, in spite of increased phosphorylation of Thr202 and Tyr204, we observed a reduction in phospho-ERK1/2 nuclear translocation and phosphorylated c-Fos. This contrasted with the effect of overexpression of Rme-6^wt^ which resulted in increased phosphorylated c-Fos. Together these results support a model whereby Rme-6 is required to ensure that, following EGFR activation, ERK1/2 is competent to translocate into the nucleus and, as discussed below, this is most likely because of Rme-6 dependent CK2 phosphorylation (Figure 6). The uncoupling of ERK1/2 nuclear translocation from the elevated ERK1/2^Thr202/Tyr204^ phosphorylation observed in the absence of Rme-6 eliminates the possibility that the effects on EGFR signalling are indirect effects on its trafficking, indicating rather that Rme-6 is playing a key informative role in regulating EGFR signalling.

**Figure 6:**
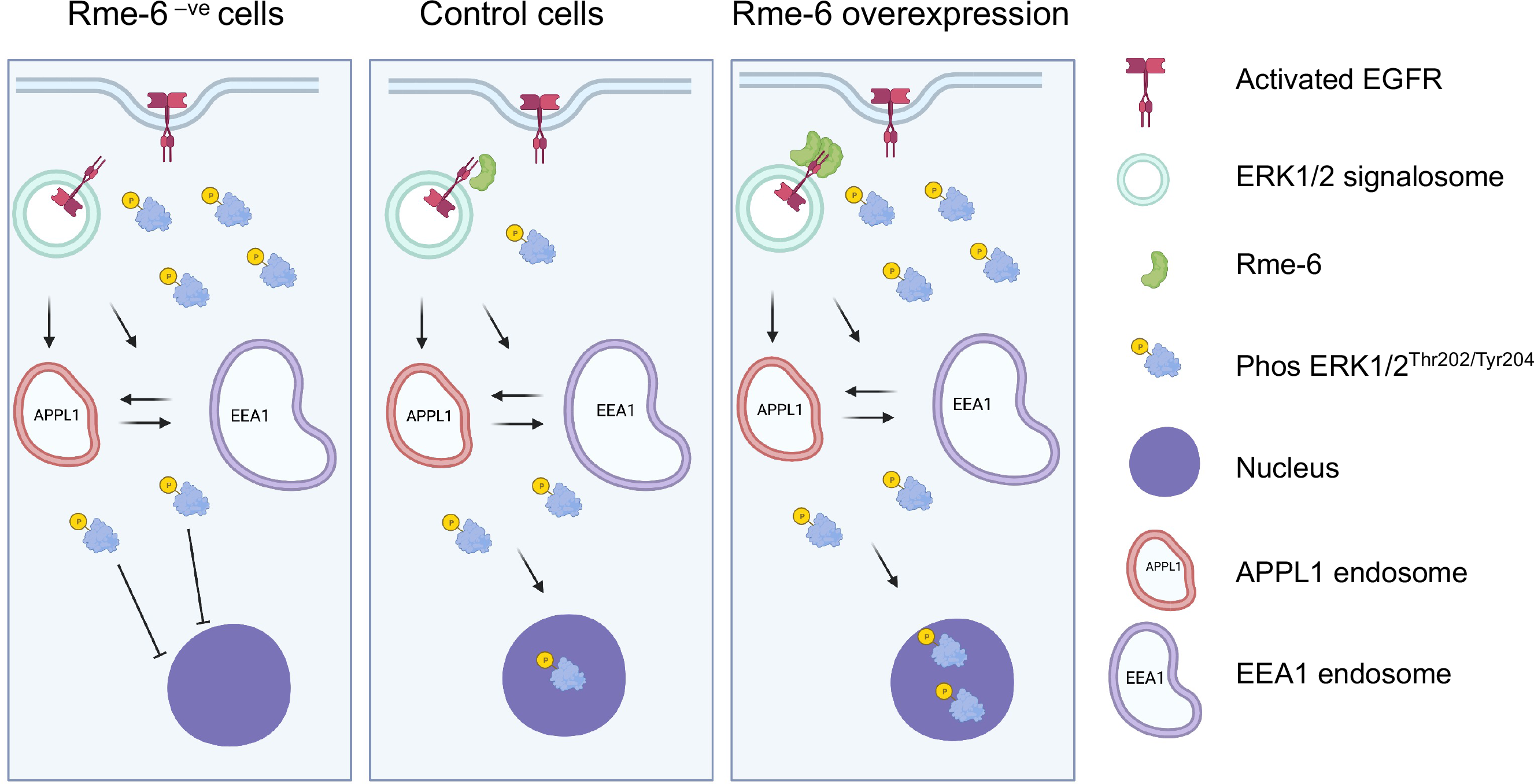
Model for an Rme-6 driven ERK1/2 signalosome. In control cells activated EGFR enters an Rme-6^+ve^ compartment competent for ERK1/2 activation (signalosome) early after internalisation. CK2 phosphorylated ERK1/2 translocates to the nucleus resulting in gene transcription. Rme-6 overexpression increases ERK1/2 nuclear translocation. Surprisingly, while loss of Rme-6 ***increases*** phosphorylated ERK1/2^Thr202/Tyr204^ in the cytoplasm, it ***reduces*** nuclear translocation of ERK1/2 and cell proliferation. Image created in BioRender.com

### Rme-6 dependent CK2 activity

CK2 is considered to be a constitutively active enzyme and it has been something of a conundrum as to how it can exert such tight control over a broad range of biological processes, which include the DNA repair response, cell proliferation, and apoptosis (Roffey and Litchfield, 2021). Our working model in this study is that the pool of CK2 that is associated with Rme-6 plays roles in EGFR signalling in two co-ordinated and complementary ways. Firstly, CK2 is required to phosphorylate ERK1/2 on Ser 244 and Ser 246 which in turn is required for effective nuclear translocation (Plotnikov et al., 2019). In the absence of Rme-6 in both HeLa Rme-6^-ve^ cells and MCF10A cells there is reduced nuclear translocation of activated ERK1/2 and reduced c-Fos phosphorylation. Our model is consistent with the pool of CK2 associated with Rme-6 ensuring effective ERK1/2 phosphorylation on Ser244 and 246. In support of specific CK2 pools that can be activated in response to individual external cues, previous studies have demonstrated that EGF can activate ERK2, resulting in increased CK2 phosphorylation of α-catenin and transactivation of β-catenin in response to Wnt signalling (Ji et al., 2009).

We have demonstrated that CK2 phosphorylation regulates the activity of Rme-6 itself. Overexpression of Rme-6^wt^ has a dominant negative effect on EGFR trafficking, delaying trafficking through APPL1- and EEA1-positive compartments. Our data show that in contrast to the overexpression of Rme-6^wt^, overexpression of Rme-6^T642D/S996D^ can rescue the effects on EGFR endocytic trafficking. Similarly, while Rme-6^wt^ overexpression results in increased nuclear phosphorylated c-Fos, overexpression of Rme-6^T642D/S996D^ results in levels of phosphorylated c-Fos which are similar to mock transfected cells. This is further supported by our results showing that purified Rme-6^T642D/S996D^ is active in the absence of any binding partners, indicating that phosphorylation at Thr642 and Ser996 sites relieves autoinhibition of Rme-6, allowing activation of the Rab5GEF activity. This is thus the second role of CK2 which ensures progression of EGFR along the endocytic pathway, limiting the activity of an EGFR dependent ERK1/2 signalosome.

### Rab5GEF driven signalosomes nuance EGFR signalling

Rme-6 is one of several Rab5GEFs which have been implicated in EGFR endocytosis and/or signalling. Rabex5 is the canonical Rab5GEF with clearly established roles in endosomal fusion and sorting into EEA1-positive endosomes (Horiuchi et al., 1997). Rin1 has a Ras binding domain and mediates Rabex5-dependent ubiquitination of endosomal Ras leading to its attenuation (Xu et al., 2010). It is striking however that loss of Rme-6 is sufficient to cause profound effects on proliferation and cell survival, indicating that there are important non-redundant functions for individual Rab5GEFs, even in the transport of a single cargo. Data from our lab and others position Rme-6 acting early after EGFR internalisation (Sato et al., 2005; Semerdjieva et al., 2008), followed by Rin1 (Balaji et al., 2012; Kong et al., 2007) and then Rabex5 (Horiuchi et al., 1997). These data are consistent with each of these Rab5GEFs establishing signalosomes that are important for a subset of EGFR signalling as has been shown in vivo (Schenck et al., 2008).

The effects of overexpression or loss of Rme-6 are relatively subtle in terms of changes in co-localisation and signalling. However, there were significant effects on both cell proliferation and apoptosis reflecting amplification of subtle changes in signalling pathways. The quite dramatic changes in cell proliferation observed following reduction in Rme-6 levels demonstrate how such subtle changes in EGFR trafficking and signalling are key to modulating the potency and the specificity of subsets of signalling (Kholodenko, 2009). It is well-established that nuclear ERK1/2 signalling is key for many cell functions and that ERK1/2 dependent gene transcription is sensitive to the duration of nuclear ERK1/2 signalling (Whitmarsh, 2011). Our studies shed important insight into EGFR dependent ERK1/2 nuclear translocation.

Rme-6 has a RasGAP domain in addition to its Rab5GEF domain and it has been shown that it has RasGAP activity *in vitro* (Su et al., 2007). However, although we observed a trend towards increased active Ras in the absence of Rme-6, we cannot be sure whether this is a direct or an indirect effect of Rme-6 RasGAP activity since other RasGAPs, e.g. p120RasGAP (Zhang et al., 2020) are present and likely to be active in HeLa and MCF10A cells.

Signalosome formation will be context-specific. In this study, for example, we showed that while Rme-6 is key for ERK1/2 signalling in HeLa cells, its loss resulted in reduced Akt signalling. By contrast reducing the levels of Rme-6 in MCF10A cells enhanced both ERK1/2 and Akt signalling, through increased protein expression, suggesting that the molecular mechanisms by which trafficking and signalling are integrated will be cell specific and indeed are likely to be context specific for particular cell types. In the latter case cross talk with other receptors and mechanical cues, among other factors, are likely to impact on the rate of flux through signalosomes. Furthermore, increasing evidence demonstrates that there is both forward and backward trafficking between endosomal populations (Flores-Rodriguez et al., 2015; Kalaidzidis et al., 2015), as well as modulation of endosome populations by signalling receptors (Villasenor et al., 2015). Together this supports a model whereby endosomal compartments are very plastic and thus can respond in sophisticated ways to extracellular signals.

In summary, our data demonstrate the role of Rme-6 in assembling and disassembling an EGFR dependent ERK1/2 signalosome. We have revealed underlying mechanisms by which a Rab5GEF co-ordinates signalling and trafficking of activated EGFR. This has important implications for understanding EGFR signalling which is often dysregulated because of mutation or overexpression in cancer. While immunotherapies have been successfully developed which inhibit aberrant signalling, the emergence of resistance is an ongoing challenge. This has led to searches for combinatorial therapies (Astsaturov et al., 2010). In this context targeting protein:protein interactions that regulate EGFR signalling is considered a promising approach (Corbi-Verge and Kim, 2016; Saafan et al., 2016). Our work therefore opens the possibility to develop novel therapeutic therapies through a molecular dissection of how EGFR flux is regulated.

## Materials and methods

### Antibodies used in this study

| Name | Host | Catalogue No. | Company | Dilution | 2° Abs | Application |
| --- | --- | --- | --- | --- | --- | --- |
| Anti Gapex5 (Rme-6) | Rabbit | A302-116A | BETHYL | 1:2000 | Alexa680 anti rabbit | WB |
| Anti-Ras cloneRAS10 | Mouse | 05-516 | MERCK | 1:1000 | LiCOR CW800 anti mouse | WB |
| Anti p44/42 MAPK(ERK1/2) L34F12 Mouse mAb | Mouse | 4696 | Cell Signaling | 1:2000 | LiCOR CW800 anti mouse | WB |
| Phospho-p44/42 MAPK (Erk1/2) (Thr202/Tyr204) (D13.14.4E) | Rabbit | 4370s | Cell Signaling | 1:2000 | Alexa 680 anti rabbit | WB |
| Anti AKT | Rabbit | 9272s | Cell Signaling | 1:1000 | Alexa 680 anti rabbit | WB |
| Anti phospho-Akt (Ser473) (587F11) | Mouse | 4051s | Cell Signaling | 1:1000 | LiCOR CW800 anti mouse | WB |
| Anti EGFR mAb(E235) | Rabbit | ab32077 | Abcam | 1:1000 | Alexa 680 anti rabbit | WB |
| Anti-GAPDH Antibody (0411): sc-47724 | Mouse | sc-47724 | Santa Cruz | 1:5000 | LiCOR CW800 anti mouse | WB |
| Anti-α-Tubulin antibody, Mouse monoclonal | Mouse | T6199 | Sigma | 1:5000 | LiCOR CW800 anti mouse | WB |
| Nucleolin | Mouse | **39-6400** | Invitrogen | 1:2000 | LiCOR | WB |
| Monoclonal Antibody (ZN004) |  |  |  |  | CW800 anti mouse |  |
| Anti EEA1 (C45B10) mAb | Rabbit | 3288 | Cell Signaling | 1:500 | Goat anti R alexa 488 | IF |
| Anti APPL1 (D83H4 mAb | Rabbit | 3858 | Cell Signaling | 1:200 | Goat anti R alexa 488 | IF |
| Anti Phospho-c-Fos (Ser32) (D82C12) | Rabbit | 5348 | Cell Signaling | 1:200<br>O/N incubation at 4C | Goat anti R Alexa 488 | IF |
| Anti Phospho-Erk1/2 (Thr202/Tyr204) | Rabbit | 9101s | Cell Signaling | 1:200 | Goat anti R Alexa 488 | IF |
| Anti-HA Cline 12CA | Mouse |  | Bioserve | 1µg/ml |  | IF |
| Anti-BrdU antibody [BU1/75 (ICR1)] - Proliferation Marker (ab6326) | Rat | ab6326 | Abcam | 1:100 | Goat Anti-Rat IgG H&L (Alexa Fluor® 647) (ab150167) | FACS |
| Goat anti-Rabbit IgG (H+L) Cross-Adsorbed Secondary Antibody, Alexa Fluor 680 | Mouse | 347580 | BD Biosciences | 1:100 | Goat anti-Mouse IgG (H+L) Alexa Fluor 488<br>#A-11017 (invitrogen) | FACS |
| Cleaved Caspase-3 (Asp175) Antibody #9661 | Rabbit | #9661 | Cell Signaling | 1:800 | Goat anti R alexa 488 | FACS |
| IRDye® 800CW | Donkey | 926-32212 | LI-COR | 1:5000 |  | WB |
| Donkey anti-Mouse IgG |  |  |  |  |  |  |

### Reagents used in this study

| Name | Catalogue No. | Company | Application |
| --- | --- | --- | --- |
| Epidermal Growth Factor, Biotinylated, complexed to Alexa Fluor™ 555 Streptavidin | E35350 | ThermoFisher | IF |
| TBB, casein kinase-2 (CK2) inhibitor | ab120988 | abcam | 2h pre-incubation |
| DAPI | 4083 | Cell Signaling | N/A |

#### Cell culture and transfections

HeLa cells were cultured in DMEM supplemented with 10% foetal calf serum, 2 mM glutamine and penicillin/streptomycin. MCF10A cells were cultured in DMEM-F-12 base supplemented with 5% horse serum, EGF (20ng/ml), recombinant insulin (10µg/ml), hydrocortisone (0.5µg/ml), 2 mM glutamine and penicillin/streptomycin. Cells were passaged every 2-3 days and were routinely tested for mycoplasma.

#### Transient transfections

Transient transfections in HeLa and Rme-6^-ve^ cells were performed using either Polyfect or Lipofectamine3000 (ThermoFisher, Ltd), according to the manufacturer’s instructions. For MCF10A siRNA mediated knockdown of Rme-6, cells were split so as to be approximately 50% confluent on the day of transfection. They were transfected with Rme-6 siRNA (GGAGAAUUCUUGAGUCGAUtt; ThermoFisher) or non-targeting siRNA (final concentration 200nM) using Lipofectamine RNAiMAX and left for 48 hours. On day 3, the cells were transfected with siRNA as before. Cells were left for 2 more days before being used for experiments. For signalling assays, cells were pre-incubated overnight in serum free medium: HeLa: DMEM containing 2mM glutamine, 1mg/ml BSA; MCF10A: complete medium without EGF or insulin.

#### Generation of Rme-6^-ve^ cells by CRISPr

HeLa cells were seeded on to 6-well plates to be 60-80% confluent on the day of transfection. Cells were transfected with guide RNAs for Rme-6 knockdown using Polyfect Transfection reagent (Qiagen), according to the manufacturer’s instructions. The guide RNA constructs were a kind gift from Horizon Discovery. The construct consisted of a gRNA sequence, Cas-9 nuclease and a GFP reporter. Positive cells were identified by GFP expression and sorted using a BDFacs Melody cell sorter and single cell cloned. Clones were verified by Western blotting.

#### Immunofluorescence

For internalization of Alexa^555^EGF, HeLa, Rme-6^-ve^ and MCF10A cells were seeded on to coverslips, serum starved overnight before being incubated with Alexa^555^EGF (5ng/ml) for indicated time points, chilled on ice and surface bound Alexa^555^EGF was acid-stripped (2 x 5 min washes with 50 mM glycine, 100 mM NaCl, 2M urea pH3, interspersed with 5 min washes with PBS/0.2%BSA). Cells were fixed with 4% paraformaldehyde (PFA). The PFA was quenched using 100mM NH_4_Cl in PBS and non-specific binding blocked with 0.2% fish skin gelatin in PBS. Cells were permeabilized using 0.1% TritonX-100/PBS. Cells were treated with primary antibodies and corresponding secondary antibodies as described in Table 1.

Coverslips were mounted onto microscopy slides and whole Z-Stacks were imaged on a Nikon wide field deconvolution microscope with a Plan Apochromat Lens 60x oil (NA 1.4) with step size of 200nm with subsequent deconvolution. Mean fluorescent intensity of internalized Alexa^555^EGF was measured in ImageJ on maximum intensity projections of Z-Stacks. Pearson’s correlation coefficient was used to measure colocalization using ImageJ. For measurement of nuclear phospho-ERK1/2 and nuclear phospho-c-Fos, overlap between antibody labelling and nuclear staining was quantified using Image J or Volocity.

#### Site directed mutagenesis

Site-directed mutations in Rme-6 were generated form a pCMV-HA-Rme-6 cDNA construct, previously described (Semerdjieva et al., 2008), using the Stratagene Quik Change mutagenesis kit according to the manufacturer’s instructions. Sequencing confirmation was performed by the University of Sheffield sequencing service.

#### Recombinant protein purification

GST-Rab5, GST-Ras and Ras binding domain (RBD)-GST were expressed and purified following expression in BL21 bacterial cells. Cells were lysed in Buffer A: 20mM Tris-HCl, pH 8.0, 500mM NaCl, 1% Triton X-100, 1 mM EDTA plus Protease cocktail (cOmplete mini, Sigma) and sonicated for 3x15sec at 14-16micron amplitude on a Soniprep150. Cell debris was pelleted at 7300 x *g* for 15 min. Following incubation of the supernatant with GSH agarose for 1 hour at 4° C, GSH agarose was washed 3 x in buffer A and recombinant proteins were eluted with buffer A containing 20mM reduced glutathione (adjusted to pH 7.5). Proteins were concentrated to ∼1mg/ml, dialysed against 20mM Tris, 100mM NaCl, snap frozen in aliquots and stored at -80°C.

Rme-6^wt^, Rme-6^T642A/S996A^ and Rme-6^T642D/S996D^ constructs were expressed using the baculovirus SF9 expression system as in (Lemaitre et al., 2019; Maib and Murray, 2022) via an N-terminal 6xHis-MBP tag. After two days of infection, the cultures were pelleted and frozen in liquid nitrogen. All pellets were thawed on ice and resuspended to 50ml final volume in 20 mM HEPES, 250 mM NaCl, 0.5 mM TCEP with the addition of protease inhibitors and Benzonase (Sigma). Cell suspensions were lysed by Dounce homogenization followed by clarification through centrifugation. Cleared lysate was filtered through 0.45 μm membranes and bound to preequilibrated Amylose Fast Flow resin (MRC-PPU Reagents and Services). After washing, protein was cleaved by addition of recombinant HRV-3C protease for at least 3 hours. Following cleavage and centrifugation, the supernatant was concentrated using Vivaspin concentrators (Cytiva) and injected onto a 24 ml Superose 6 Increase 10/300 GL equilibrated in 20 mM HEPES, 250 mM NaCl, 0.5 mM TCEP. Peak fractions were analyzed by SDS-PAGE, pooled and concentrated and snap frozen in aliquots.

#### Rab5GEF assays

GEF assays were performed as described (Kulasekaran et al., 2021). Briefly GST-Rab5, purified from BL-21 bacteria by affinity chromatography, was loaded with GDP by incubating for 20 min at 30°C in 20mM Tris, pH 7.5, 100mM NaCl and 5mM EDTA containing 20µM GDP. Rab5GDP was stabilised by the addition of 10mM MgCl2. Exchange reactions contained 0.2µM preloaded active Rab5, and 0.5-3 µM Rme-6 wild type and mutants, in 20mM Tris, pH 7.5, 100 mM NaCl, 0.5mg/ml BSA, 0.5mM dithiothreitol and 5µM α^32P^GTP (specific activity: 5µCi per nmole, Perkin Elmer). Exchange reactions were incubated at 30°C for 30 minutes before being quenched with 1 ml of ice cold 20mM Tris, pH 7.5, 100 mM NaCl and 20mM MgCl_2_ (wash buffer). Samples were applied to nitrocellulose membranes under suction and membranes were washed with 5ml of ice cold wash buffer before being counted using Cerenkov counting in a Wallac scintillation counter.

#### In vitro kinase assays

In vitro kinase assays were performed in 20 mM Tris, pH 7.2, 100mM NaCl and 1 mM MgCl_2_ using 6 nmoles Rme-6 wild-type and mutants, CK2 (500 units, NEB Ltd) and 100µM ^32P-^ATP (specific activity: 1µCi per nmole, Perkin Elmer). Kinase assays were performed at 30 °C for 30 minutes and were terminated by addition of Laemmli sample buffer before SDS-PAGE and exposure to film. Bands were quantified using ImageJ.

#### Measurement of active Ras

Active Ras was measured using the Ras binding domain of Raf (RBD) (Baker and Rubio, 2021). HeLa and MCF10A cells were serum starved overnight in serum free medium (DMEM containing 1mg/ml BSA). On the day of the experiment cells were incubated with 5ng/ml EGF for various times. Cells were chilled on ice and washed 3x with ice cold PBS. Cells were lysed in buffer B: 50 mM Tris-HCl, pH 7, 10% Glycerol, 200 mM NaCl, 2.5 mM MgCl_2._

Following incubation for 15 minutes on ice, cell lysates were centrifuged for 15 minutes at 4°C at 20,000 x *g*. Equal amounts of protein were then loaded onto RBD-GST bound to GSH agarose. Samples were incubated at 4°C for 1 hour with rolling. Beads were washed 3x in lysis buffer and proteins eluted by addition of Laemmli sample buffer and boiling for 5 minutes at 95°C. Samples were then subjected to SDS-PAGE and Western blotting, probing with anti-Ras antibodies.

#### BrdU proliferation assay

MCF10A cells which had been treated with either control siRNA or siRNA targeting Rme-6 were seeded at a density of 23-30% and left for 23.5 hours. BrdU (final concentration:25µM) was added to cells for I hour. Media was removed and the cells were washed once in PBS. Cells were removed from the plate by trypsinisation, diluted in PBS and centrifuged at 500 x g for 5 min. Cells were resuspended in 0.5ml PBS + 2mM EDTA. 4.5ml ice-cold 70% ethanol was added dropwise, and cells were chilled at 4°C until ready to use (overnight at least) or kept at -20°C for long storage. Cells were centrifuged at 1000 x *g* for 5 mins to pellet cells and the supernatant was discarded. Cells were resuspended in1ml 2M HCl, and incubated at room temperature for 20min. Cells were washed once in PBS/2mM EDTA and then resuspended in 0.1M sodium borate, pH8.5 (0.5ml). Following incubation at room temperature for 3 min cells were washed 1x and resuspended in 0.1ml dilution buffer (0.5% BSA + 0.2% Tween in PBS) and primary antibodies added for 2 hours at 4°C with gentle agitation. Cells were washed 3 x and incubated with secondary antibodies for I h at 4°C. Following 3 washes, cells were resuspended in 1ml PBS + 2mM EDTA containing 20ug/ml propidium iodide and 200ug/ml RNAse A (Code#EM101-02-01, Lot#M31122, 10mg/ml) and analysed by FACS using an LSRII Facs.

## Acknowledgements

Work was funded by the Wellcome Trust: 087639/Z/08/Z (E.S), 211193/Z/18/Z (D.H.M), an MRC studentship (S.S), the University of Bisha, Saudi Arabia (F.A), a University of Sheffield Centre for Membrane Interactions and Dynamics studentship (Z.Z.). Fluorescence microscopy was carried out in the Wolfson Light Microscopy Facility funded by MRC grant (MR/K015753). We thank Svetlana Sedelnikova for preparation of GST-Rab5.

