## Supplementary figures and images for "Rme-6 integrates EGFR trafficking and signalling to regulate ERK1/2 signalosome dynamics"

### Supplementary Figures

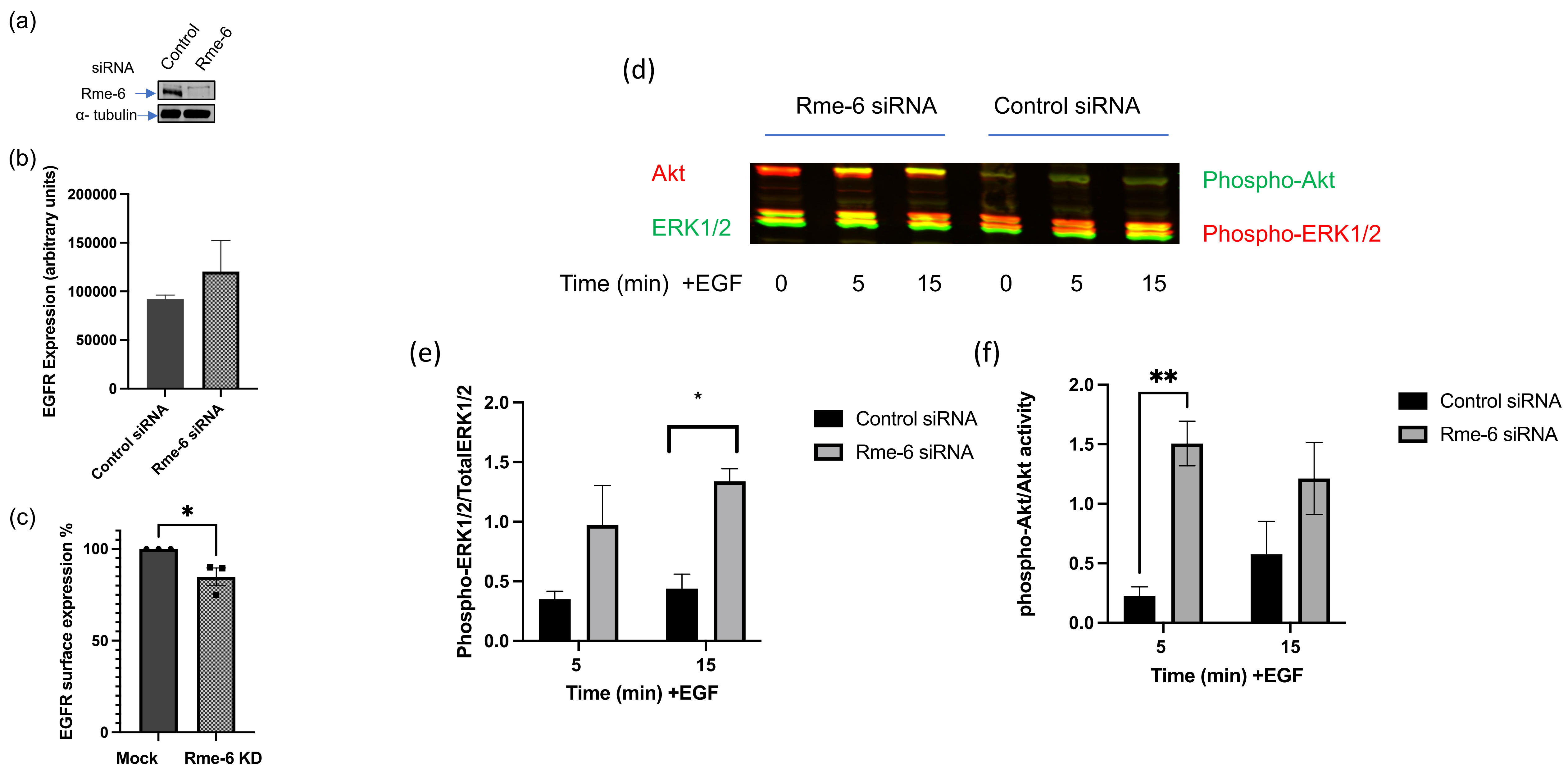


Supplemental Figure 1

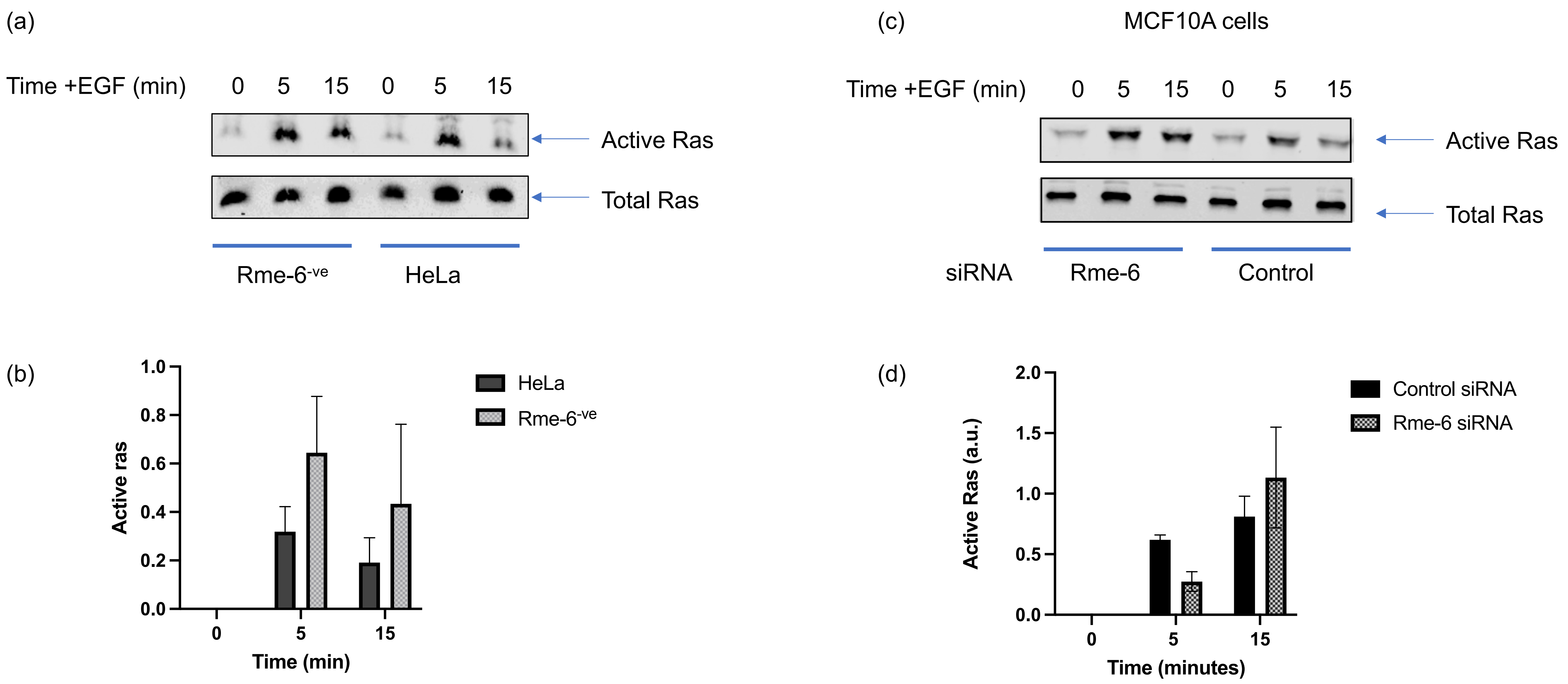


Supplementary figure 2

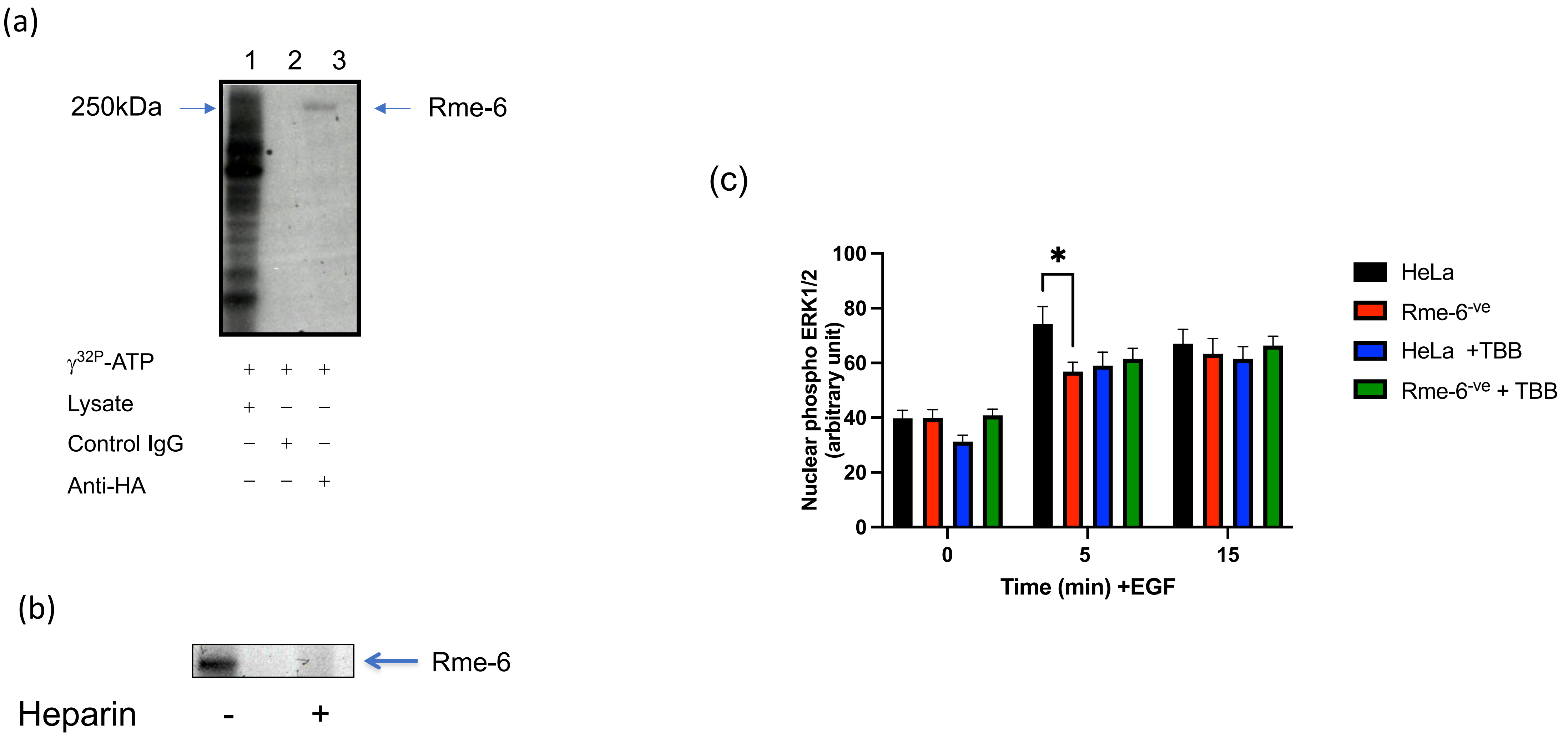


Supplementary Figure 3
