## Supplementary Legends for "Rme-6 integrates EGFR trafficking and signalling to regulate ERK1/2 signalosome dynamics"

Supplementary Figure Legends

*Supplementary Figure 1: Loss of Rme-6 affects EGFR expression and signalling in MCF10A cells*

(a) Western blots of lysates from MCF10A cells treated with control siRNA or siRNA targeting Rme-6. Blots were probed with antibodies against Rme-6 and α-Tubulin as a loading control.

(b) Expression of EGFR in MCF10A cells treated with control siRNA or siRNA targeting Rme-6 was measured by Western blotting. Data are the mean +/-SEM of three independent experiments.

(c) Surface expression of EGFR on MCF10A cells treated with control siRNA or siRNA targeting Rme-6. Data are the mean +/-SEM of three independent experiments. *:p<0.05

(d) Representative Western blot showing total ERK1/2 (green) and phosphorylated ERK1/2 (red) in MCF10A cells treated with control siRNA or siRNA targeting Rme-6 following EGF stimulation (5ng/ml) for the indicated times.

(e) Quantitation of the ratio of phospho-ERK1/2 to total ERK1/2 from three independent experiments. Data are expressed as mean +/- SEM. *<0.05.

(f) Quantitation of the ratio of phosphorylated Akt to total Akt from three independent experiments. Data are expressed as mean +/- SEM. **: p<0.01

*Supplementary Figure 2: Loss of Rme-6 increases active Ras in Hela and MCF10A cells*

Active Ras was captured by RBD-GST from cell lysates following treatment with EGF (5ng/ml) for the indicated times. Representative Western blots showing total and active Ras from HeLa and Rme-6$^{-ve}$ cells (a) and MCF10A cells treated either with control siRNA or siRNA targeting Rme-6 (b). Quantitation of active Ras from three independent experiments in HeLa and Rme-6$^{-ve}$ cells (c) and MCF10A cells treated either with control siRNA or siRNA targeting Rme-6 (d). Data are expressed as mean +/-SEM of three independent experiments.

*Supplementary Figure 3: A pool of CK2 is associated with Rme-6 and is required for ERK1/2 nuclear translocation*

(a) Lysates (lane 1) from Hek293T cells transfected with HA-Rme-6 were prepared and incubated with Protein G beads coupled to control IgG (lane 2) or anti-HA antibodies (lane 3). Beads and lysate were then incubated with $\gamma$32P-ATP for 30 min at 30°C before SDS-PAGE and autoradiography. In the absence of added kinases, immunoprecipitated Rme-6 is phosphorylated.

(b) HA-Rme-6 which had coupled to Protein G beads was incubated in the presence or absence of 0.5µg/ml heparin, a known inhibitor of CK2 (Hathaway et al., 1980). In the presence of heparin phosphorylation is significantly reduced suggesting that CK2 is closely associated with HA-Rme-6.

(c) Quantitation of nuclear localisation of phospho-ERK1/2 in HeLa and Rme-6$^{-ve}$ cells in the presence or absence of the CK2 inhibitor, (4,5,6,7-tetrabromobenzotriazole, TBB) in three independent experiments. Data are expressed as mean +/-SEM. Significance was determined with a Students T-Test. *: $p<0.05$;
